# Flexibility and Neural Correlates of Action-Sound Predictions

**DOI:** 10.1101/2025.03.19.644065

**Authors:** Fabian Aurich, Andreas Widmann, Tjerk T. Dercksen, Betina Korka, Anni Richter, Max-Philipp Stenner, Nicole Wetzel

## Abstract

To interact efficiently with our environment, our brain predicts the sensory effects of our actions and compares them with the actual outcomes. This allows us to adapt our actions when predictions and sensory outcomes mismatch. While this process is generally well understood for action-sound predictions, it is an open question how flexibly these predictions can adapt in frequently changing environments, as they occur in real life.

To investigate the flexibility of top-down predictions, we asked participants (N = 41) to press one of two buttons, a left-hand and a right-hand button, and switch hands autonomously. One button frequently produced a sound (80%) and rarely no sound. The other button frequently generated no sound (80%) and rarely produced a sound. In a third, separate condition, each button produced a sound in 50% of the trials.

Unexpected sounds and unexpected sound omissions elicited a series of error-related brain responses in the electroencephalogram (EEG) at different levels of auditory processing, including a mismatch negativity (MMN) and the P3 complex for unexpected sounds, and the oN1, oN2, and oP3 complex for unexpected omissions. Moreover, unexpected sounds elicited an equivalent MMN—regardless of whether silence was expected (80%) or no reliable expectation was possible (50%), while later P3 components showed different amplitudes.

Our results demonstrate flexible action-sound predictions at sensory and higher cortical levels. Furthermore, they indicate that predicted silence does not have an explicit sensory representation at lower levels but emerges at later stages, when higher-level information has been integrated.

## 1. Introduction

Our actions often lead to expectable sensory outcomes, which can help guide our behaviour. Imagine typing a message on your smartphone: each keystroke triggers an expected subtle “click” sound, giving you feedback that the input has been successful. In case of a missing feedback, we are surprised and double check (e.g., Baragona et al., 2025). Contrary, when you silence your phone by putting it in “mute” mode, you expect silence (the absence of auditory feedback) after a keystroke and may be surprised otherwise. Each individual action results in a specific action effect. Consequently, expectations of action-effects must be flexible, depending on action and context. This study examines the generation of flexible action-based auditory predictions and the brain mechanisms involved. To this end, we violated auditory predictions in two ways, by presenting a sound when no sound was expected, and by omitting an expected sound. We compared the elicited brain responses to physically identical events when they were expected vs. unexpected. Furthermore, we examined the brain responses to the same events when no clear prediction can be formed, such as when typing on a new, unfamiliar phone.

Predictive coding suggests that higher and lower cortical levels of the brain communicate in a continuous loop to predict future events of our surroundings. In this loop, sensory predictions generated at higher cortical levels are sent down to lower levels (Clark, 2013; Feldman & Friston, 2010). At lower levels, sensory inputs received from the environment are compared with the top-down sensory predictions (Arnal & Giraud, 2012; Clark, 2013; Friston, 2010). When mismatches occur, prediction errors are generated. These error signals can be sent back to higher levels to serve as feedback to refine future sensory predictions and minimize prediction errors (Feldman & Friston, 2010).

An important source of predictions are our own actions, given that much of the sensory input we receive is self-generated and therefore predictable. For example, typing on your phone generates a predictable sound. By comparing predicted and actual sensory input you can verify whether the motor action fulfilled the intention to produce the correct input on your phone. Prediction is therefore central to theories of motor control. This includes ideomotor theory, which suggests that actions are guided by the expected sensory effects they aim to produce (Greenwald, 1970; Hommel, 2003, 2013). Initially, through repeated associations of motor actions with subsequent sensory inputs, a strong link between action and sensory effect develops. This is known as ideomotor learning, a prerequisite for motor control mechanisms to work. Once these action-effect associations are established, we choose and evaluate our actions based on their desired effects and intention (Elsner & Hommel, 2001). The Theory of Event Coding (TEC) extends this by proposing that actions and their sensory effects are neurally encoded as integrated representations, or *event files*, which are retrieved based on the agent’s current goals (Hommel, 2019; Hommel et al., 2001).

Building on ideomotor theory, the *extended Auditory Event Representation System* (eAERS) described by Korka et al. (2022) integrates sensory and ideomotor information into a unified sensory predictive model at lower cortical levels to explain how the brain processes and predicts auditory stimuli. This predictive model combines information from bottom-up sensory regularities with top-down ideomotor expectations to form predictions about future auditory input. New incoming auditory input is then compared with these predictions and any mismatches elicit prediction error signals. These signals, along with auditory and relevant non-auditory information, are integrated to form auditory event representations and are propagated to predictive models in higher-level cognitive systems for further evaluation to guide attention allocation and refine motor behaviour.

Previous research has primarily employed static paradigms, focusing on single action-effect contingencies across blocks of trials (Dercksen et al., 2024; Dercksen et al., 2020; Nittono, 2006; SanMiguel, Widmann, et al., 2013). These paradigms fail to capture the dynamic flexibility required in real-world contexts, where actions and their effects frequently change. This raises the question of whether action-based auditory predictions are dynamically adjusted when actions and contexts frequently change.

The present event-related potential (ERP) study addresses this knowledge gap, by investigating whether auditory top-down predictions based on self-produced actions can be changed on a trial-by-trial basis. We therefore employed a bimanual button press paradigm to establish multiple action-effect contingencies simultaneously. For instance, Korka et al. (2019) examined action-intention violations using left and right button presses to generate high and low frequency standard tones, with occasional unexpected deviant tones. They found that early auditory ERPs, such as the Mismatch Negativity (MMN) and the P3a component, were elicited when participants’ top-down expectations, based on their intended actions, were contradicted (see also Widmann & Schröger, 2022). The MMN, an early pre-attentive response to deviations from an established auditory pattern (Näätänen et al., 2007), is discussed to represent the mismatch between the predictive model and sensory representations (Garrido et al., 2009) in action-based (Korka et al., 2019; Widmann & Schröger, 2022) but also regularity-based paradigms in auditory (Näätänen, 1990; Näätänen et al., 2007), visual (Pazo-Alvarez et al., 2003) and somatosensory (Näätänen, 2009) domains. The MMN is often followed by a P3a component. The P3a component indicates an involuntary attention shift towards the deviant stimulus (Escera et al., 1998; Waszak & Herwig, 2007), depending on its relevance to the listener’s current goal (István Winkler & Schröger, 2015). It has been shown that the P3a, like the MMN, is also sensitive to violations of action-based predictions (Korka et al., 2019; Nittono, 2006; Widmann & Schröger, 2022). This component is thus considered as the next stage in auditory distraction processing (Horváth et al., 2008). It is part of a broader P300 complex, including P3b and the Novelty P3 subcomponents (Barry & Rushby, 2006; Barry et al., 2020) that presumably reflect higher cognitive evaluation tasks in auditory processing (Polich, 2007) along the eAERS hierarchy (Korka et al., 2022). This makes the P300 complex, alongside the MMN, suitable for the current study to investigate the flexibility of intention-based top-down prediction control.

However, a challenge in investigating auditory prediction error-related mechanisms is the clear differentiation between bottom-up and top-down effects on the generation of prediction errors: “The informational content of the prediction error is in fact composed of two parts: that part of the sensory input that is encountered but was not predicted [bottom-up], and that part of the sensory input that was predicted but is in fact absent [top-down]” (Schröger et al., 2015, p. 647). Thus, the present study aims to separate the two types of prediction violation typically occurring concurrently: (a) a stimulus which is not expected but presented (i.e., mispredicted sound) and (b) another stimulus which is expected but not presented (i.e., mispredicted sound omission). The latter type of violation (b) hereby minimises the influence of bottom-up signals on the measured indicators of the prediction errors. Thus, the measured neural response in this case reflects the mismatch between the top-down prediction of an auditory action effect and the unexpected absence of auditory input. It has been discussed that the brain’s mismatch response in the time range of the N1 component represents this prediction error (’omission N1’– oN1), as there is no interfering sound information to overlay the prediction error signal (SanMiguel, Widmann, et al., 2013; Wacongne et al., 2011). Moreover, an increasing number of auditory omission studies investigated these prediction errors in humans and reported a sequence of brain omission responses consisting of oN1, oN2 and oP3 (Dercksen et al., 2020; SanMiguel, Saupe, & Schröger, 2013; SanMiguel, Widmann, et al., 2013; van Laarhoven et al., 2017).

In sum, this study investigates whether and how flexible action-based auditory predictions are generated on a trial-by-trial basis. Action-based predictions were violated by the presentation of an unexpected sound or the unexpected omission of a sound. We asked participants to press one of two buttons with their left or right hand and change the hand from trial to trial. One button frequently produced a sound and rarely omitted the sound. The other button frequently generated no sound and rarely produced a sound. If flexible top-down predictions are established on a trial-by-trial basis (changing the hand pressing the button), the violation of predictions is expected to evoke MMN and P3 components when an unexpected sound is presented, and oN1, oN2 and oP3 components when a sound is unexpectedly omitted. Moreover, in a third condition sounds were presented with a probability of 50% for each hand, so that participants could not form a reliable prediction whether, or not, a sound will be generated (SanMiguel, Widmann, et al., 2013; Tast et al., 2024).

## 2. Materials and Methods

### Participants

A power analysis using GPower (Faul et al., 2009) determined that a minimum of 34 participants was required for analysis to achieve a power of 0.8 for the planned paired-samples *t*-tests (*Cohen’s d* = 0.5, *α* = 0.05) and 28 participants for the repeated measures ANOVA (*f* = 0.25, *α* = 0.05). To ensure a robust power across analyses, we aimed for a larger sample size. Therefore, 41 participants (31 female, 10 male) with a mean age of 25.3 years (range: 18– 40, *SD* = 4.6) took part. Handedness, assessed via an adapted German Oldfield Scale (Oldfield, 1971), identified 40 right-handed participants and 1 left-handed participant. Recruitment involved posters and social media. Exclusion criteria were an age outside 18-40, impaired hearing or diagnosed neurological or psychiatric disorders stated by participants. All participants signed written informed consent before the experiment. Ethical approval from the Ethics Committee of the Medical Faculty of the Otto-von-Guericke University Magdeburg was obtained via an amendment to the project 189/18. Participants received either monetary compensation or course credits for participation.

### Stimulus and Apparatus

Participants were seated on a chair at a table in a dimly lit booth that provided both acoustic and electromagnetic shielding. The experiment was created using Psychtoolbox (Brainard, 1997) and ran on a Linux-based system with GNU Octave (version 4.0.0.). Participants responded via custom-built infrared photoelectric buttons, one for each index finger. The buttons were covered with sound-absorbing soft foam to minimize any button press related noise, as used in previous auditory omission studies (Dercksen et al., 2020; Korka et al., 2020). The buttons were linked to the stimulus computer via an RTbox (Li et al., 2010).

Visual stimuli were presented on a VIEWPixx/EEG Display (Resolution 1920(H) x 1080(V) – 23.6-inch display size), approximately 60 cm away from the participants. The stimuli included a white fixation cross (0.67° x 0.67° visual angle) displayed at the centre of a grey background (RGB: 128, 128, 128) and additional versions of a speaker symbol (2.38° x 2.38° visual angle) or question mark (1.52° x 1.39° visual angle) symbols symmetrically placed on both sides of the cross (**Figure 1**). Binaural auditory stimuli were delivered simultaneously via two loudspeakers (Bose Companion 2 series III Multimedia speaker system) located on the left and right sides of the screen. The stimuli comprised a 100 ms, 1000 Hz low-pass filtered (fourth-order Butterworth filter; SanMiguel, Widmann, et al., 2013), click sound with a sound pressure level of 72 dB. The loudness was held constant for all participants.

**Figure 1.**
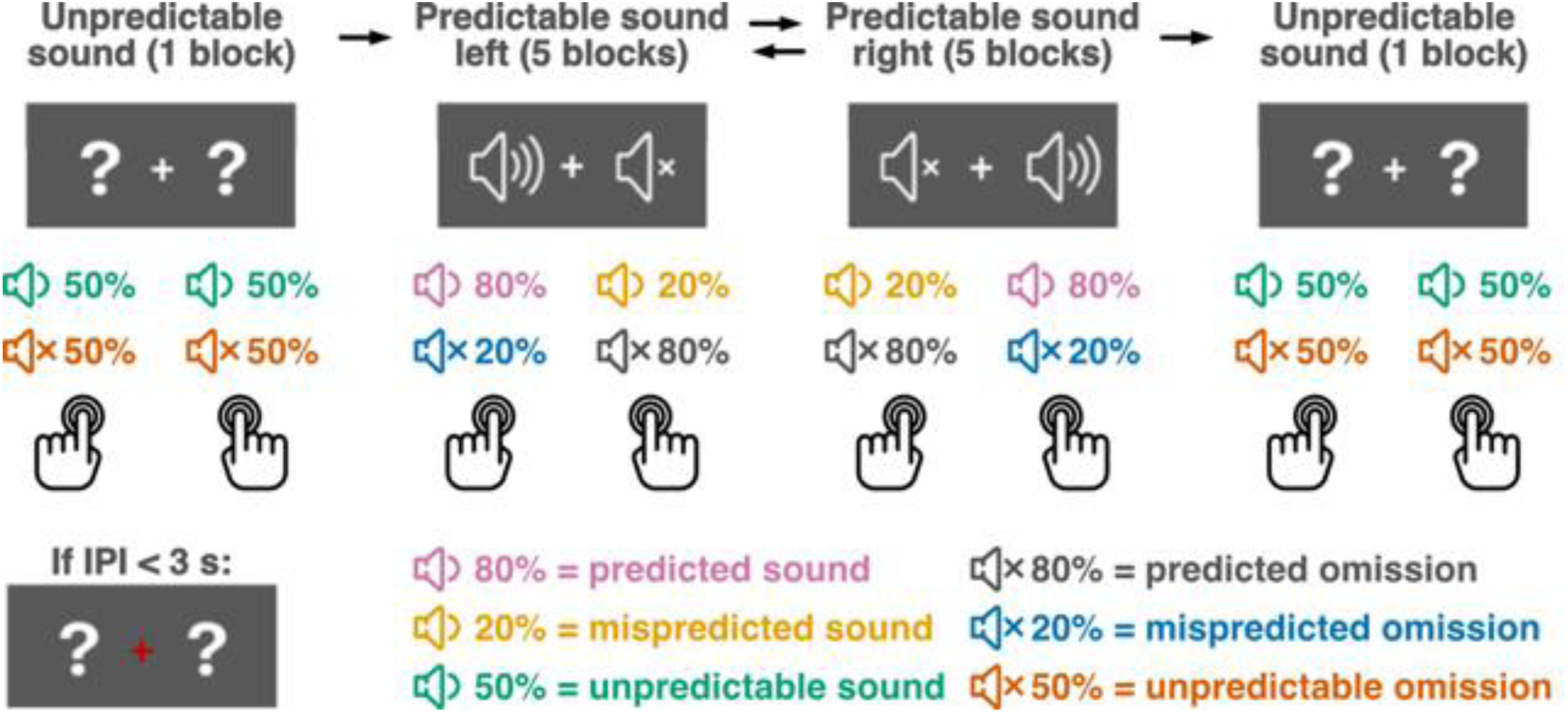
Schematic description of the experimental design. A total of twelve blocks were conducted successively. The initial block was always an Unpredictable Sound block, followed by five Predictable Sound Left blocks, five Predictable Sound Right blocks and a concluding Unpredictable Sound block. The order of the Predictable Sound Right and Predictable Sound Left blocks was counterbalanced across participants. In the Predictable Sound Left blocks, the predicted sound was presented on the left side in 80% of the button presses, whereas the mispredicted sound was presented on the right side in 20% of the button presses. The opposite was the case during the Predictable Sound Right blocks.

### Task and Procedure

Participants carried out self-paced button presses with their left and right index fingers. Each finger had a separate button. For any given press, participants decided for themselves which button to press. However, we asked participants to press each of the two buttons approximately equally often within any given block (of 90 presses each), and to refrain from any deliberate, fixed patterns in their choice (e.g., alternating between buttons, or higher-level patterns).

Participants had to wait for at least 3 seconds between consecutive button presses (inter-press interval, IPI). If they pressed earlier than 3 seconds after the preceding press, the fixation cross turned red for 500 ms. After each block, we informed participants about the number of button presses that had been too early and the average IPI for that block.

Pressing a button generated an immediate click sound with a certain probability, which could differ between the two buttons depending on the block type. There were three block types. In *Predictable Sound Left* blocks, pressing the left button generated a sound in 80%, and no sound in 20%, while pressing the right button generated a sound in 20%, and no sound in 80% of cases (**Figure 1**). In *Predictable Sound Right* blocks, the right button generated a sound in 80%, and no sound in 20%, while the left button generated a sound in 20%, and no sound in 80% of cases. In *Unpredictable Sound* blocks, each button generated a sound in 50%, and no sound in 50% of cases.

Across the three block types, this design yielded six event types. When a participant pressed a button that was associated with an 80% sound probability and actually generated a sound, this event defined a *predicted sound* trial. Generating no sound after pressing a button that was associated with an 80% sound probability defined a *mispredicted omission* trial. When a participant pressed a button that was associated with a 20% sound probability and generated no sound, that event defined *predicted omission* trials, while generating a sound in response to that button press defined *mispredicted sound* trials. Finally, generating a sound, or no sound, in *Unpredictable Sound* blocks defined *unpredictable sound* trials and *unpredictable omission* trials, respectively.

Throughout the *Predictable Sound Left* and *Predictable Sound Right* blocks, participants saw a speaker symbol on one side of the fixation cross and a muted speaker symbol on the other side, illustrating the probability of sound presentation for each button. This reinforced the unbalanced probability distribution of a *predicted* (higher probability of presentation) and a *mispredicted* (lower probability of presentation) *sound* following a button press. In the *Unpredictable Sound* blocks, two question marks were positioned on either side of the fixation cross (**Figure 1**).

The experiment consisted of 12 blocks. All participants first completed an initial *Unpredictable Sound* block, followed by five consecutive blocks each for the *Predictable Sound Left* and *Predictable Sound Right* conditions, and a final *Unpredictable Sound* block. The *Unpredictable Sound* blocks were split in order to balance effects of attentional state, practice, and fatigue as much as possible (as done similarly for control conditions in preceding studies; e.g., Baragona et al., 2025; Dercksen et al., 2020; SanMiguel, Saupe, & Schröger, 2013; SanMiguel, Widmann, et al., 2013). The order of the *Predictable Sound Left* and *Predictable Sound Right* conditions was counterbalanced across participants. Before the main experiment, all participants completed one training block of each block type, to get attuned to the task and the pace of button presses. Training started with a *Predictable Sound Left* followed by a *Predictable Sound Right* block and ended with an *Unpredictable Sound* block. Training blocks were shorter than experimental blocks and consisted of 30 trials each. Participants received feedback about the number of button presses that had been too early and the average IPI after each training block.

We defined each button press as a trial. Each block ideally consisted of 90 trials, resulting in a total of 1080 trials. For each block, the presentation of a sound after a given button press was pseudo-randomized. The *Predictable Sound Left* and *Predictable Sound Right* blocks were designed to include 450 trials each, ideally comprising 360 predicted trials (180 sound and 180 omission trials) and 90 unpredicted trials (45 sound and 45 omission trials). In the *Unpredictable Sound* block, participants completed 180 trials, with a target distribution of 90 unpredictable sound and 90 omission trials. The first five trials of every *Predictable Sound Left* / *Predictable Sound Right* block were always predicted events. Following each mispredicted event in the *Predictable Sound Left* / *Predictable Sound Right* block, two consecutive predicted events followed. Although these numbers reflect the ideal event distributions, actual trial counts were likely to vary across participants due to differences in button press behaviour, such as unequal left and right button press patterns. To minimize these deviations and maintain the target ratios, the stimulus presentation (sound or sound omission) was dynamically adjusted based on the monitored history of participants’ button presses. This trial-adaptive approach ensured that the target ratios were approximated.

### EEG recording

We recorded from 63 active electrodes placed according to the extended International 10-20 system. The active electrodes were placed at Fp1/Fp2, AFz, AF3/4, AF7/8, Fz, F1/2, F3/4, F5/6, F7/8, FC1/2, FC3/4, FC5/6, FT7/8, Cz, C1/2, C3/4, C5/6, T7/8, CPz, CP1/2, CP3/4, CP5/6, TP7/8, Pz, P1/2, P3/4, P5/6, P7/8, POz, PO3/4, PO7/8, O1/2 and at the left and right mastoids (M1/M2). Electrooculography (EOG) electrodes were positioned at the lateral canthi of both eyes and below the left eye. A reference electrode was placed at the tip of the nose. We recorded via an Actichamp amplifier (BrainProducts, Gilching, Germany) at a sampling frequency of 500 Hz using the Vision Recorder software (version 1.21).

### Data analysis

**EEG data preprocessing** and analysis were performed using the MATLAB software (version R2022b) and the EEGLAB toolbox (Delorme & Makeig, 2004). Two datasets were prepared. For the analysis dataset, the raw EEG data was filtered using a 0.1 Hz high-pass filter (Kaiser windowed sinc FIR filter, order = 8024, beta = 5, transition bandwidth = 0.2 Hz) and a 48 Hz low-pass filter (Kaiser windowed sinc FIR filter, order = 402, beta = 5, transition bandwidth = 4 Hz). The data was then divided into epochs, with each epoch covering a period from 200 ms before the button press to 500 ms after. Trials were excluded if the IPI fell below 3000 ms, or above 8000 ms. Channels with a robust z-score of the robust standard deviation of their amplitudes over time larger than three were removed (Bigdely-Shamlo et al., 2015). Epochs that exceeded a 500 µV signal change threshold were discarded. A second dataset was derived from the raw data to facilitate detection of artifacts associated with heart– and eye-related activity using independent component analysis (ICA; Makeig et al., 1995). This dataset was identical to the analysis dataset (the same trials and channels were removed), with the difference that the raw data was high-pass filtered at 1 Hz in order to improve ICA decomposition (Klug & Gramann, 2021; Irene Winkler et al., 2015), using a Kaiser windowed sinc FIR filter (order = 1604, beta = 5, transition bandwidth = 1 Hz), followed by a low-pass filter at 48 Hz using a Kaiser windowed sinc FIR filter (order = 402, beta = 5, transition bandwidth = 4 Hz). After ICA was performed on this dataset, the obtained demixing matrix was applied to the 0.1–48 Hz filtered data, which was then used for further analysis.

ICs were classified with the support of the IClabel plugin (Pion-Tonachini et al., 2019). Two independent evaluators (FA and TD) visually inspected the components for heart– and eye-related artifacts, and discussed the components that each evaluator had identified as artifacts to reach a consensus. These components were then removed from the dataset. Subsequently, channels that had been identified as bad channels, and removed, were interpolated. Trials exceeding a 125µV signal change threshold were excluded. Epochs were baseline corrected by subtracting the mean amplitude within the time interval between 200 ms and 100 ms prior to stimulus onset (to exclude motor execution potentials, as done similarly in previous studies, e.g., Dercksen et al., 2024; Dercksen et al., 2020). The initial five trials of each block, and the two trials following a mispredicted event, which were always predicted events, were excluded (Dercksen et al., 2024; Dercksen et al., 2020).

Lastly, we averaged data across trials to obtain an event-related potential (ERP) for every participant and condition. Due to the bimanual design of the experiment, contralateral and ipsilateral events were recorded on electrodes of both hemispheres. For analysis, all contralateral data were merged onto the corresponding electrodes of the right hemisphere, and all ipsilateral data onto the corresponding electrodes of left hemisphere, as illustrated in **Figure 2**.

**Figure 2.**
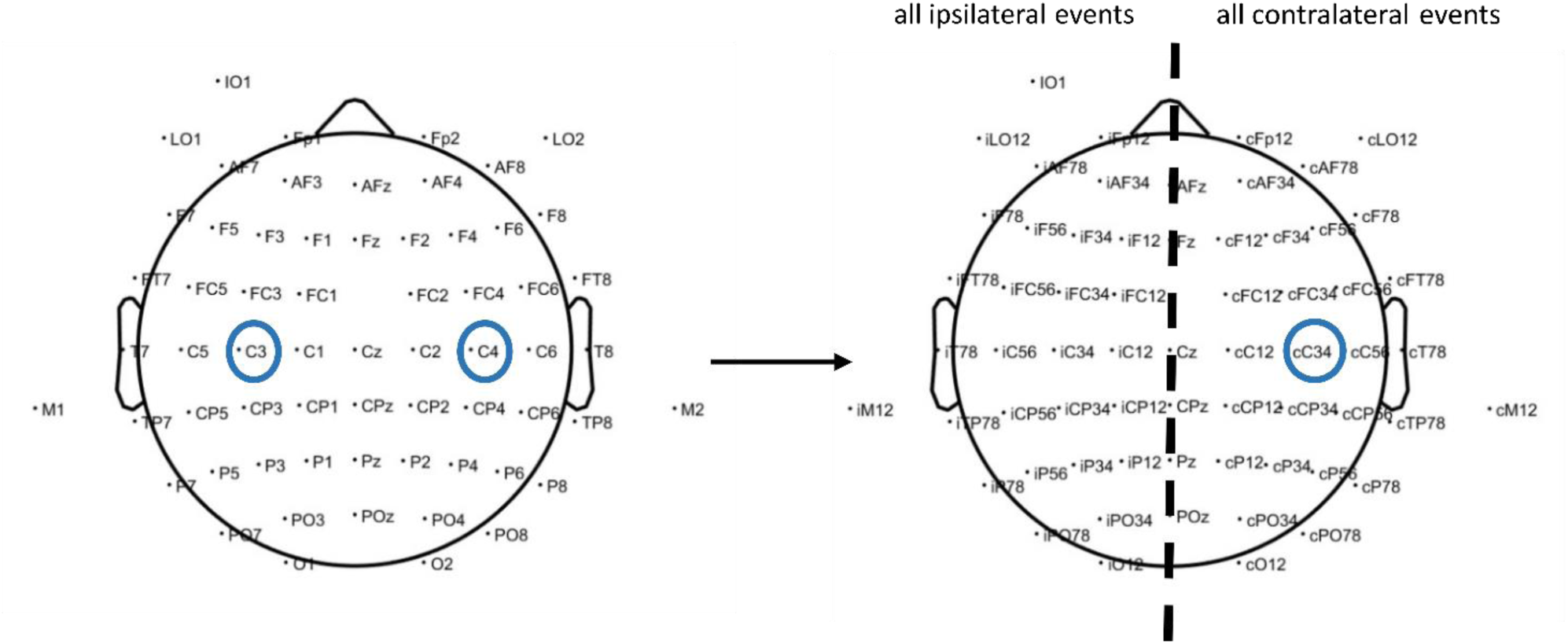
Re-definition of channels as ipsilateral or contralateral to movement. When collapsing across trials, channels contralateral to movement were distributed across both hemispheres because participants moved the left and right index finger in different trials. To disentangle contralateral from ipsilateral signals, we defined contralateral channels (denoted with a small c, e.g., cC34) by collapsing data from left-hemisphere channels during right-hand movements (e.g., C3), and from right-hemisphere channels during left-hand movements (e.g., C4). Throughout the manuscript, contralateral channels are shown over the right hemisphere, and ipsilateral channels over the left hemisphere (denoted with a small i, e.g., iC34).

### Behavioural data

Participants were instructed to press the left and right buttons approximately equally often during all block types (*Predictable Sound Left, Predictable Sound Right, Unpredictable Sound*) to ensure a consistent probability distribution of sound presentation for both hands. This way, auditory adaptation, occurring in response to repeated sound, should remain constant throughout the experiment (i.e., 50% sound probability across trials in all conditions). To verify that the frequency of sound presentation was indeed balanced across block types, we calculated the ratio of left to right button presses for each block type. Furthermore, we compared median inter-press intervals between block types after removing button presses with an IPI less than 3000 ms, or greater than 8000 ms.

### PCA

Temporal Principal Component Analysis (PCA) was performed to investigate sound– and omission-related ERPs, by decomposing them into individual components that collectively contribute to shaping the resulting ERP waveforms (Scharf et al., 2022). The number of components to be retained was determined using the Empirical Kaiser Criterion (Braeken & van Assen, 2017; Scharf et al., 2022). A Geomin rotation (ε = 0.01), implemented in R (R 4.1.2; R Core Team, 2021), was applied to the initial PCA solution as described in the tutorial of Scharf et al. (2022).

Two separate PCAs were conducted in order to consider a potential different structure of components evoked by sound and omission trials (Dercksen et al., 2020). The sound PCA focused on the analysis of auditory stimulus responses and included per participant averaged ERPs for *predicted sounds*, *mispredicted sounds*, and *unpredictable sounds*. A second omission PCA focused on the omitted auditory stimulus responses and included per participant averaged ERPs for *predicted omissions*, *mispredicted omissions*, and *unpredictable omissions*.

We selected channels for statistical analysis by visual inspection of the topography of PCA components of interest. Regarding the sound components, the following channels of interest were selected: Fronto-central channels (Cz, Fz, iFC1/2, cFC1/2) for MMN and P3a, centro-parietal channels (Cz, CPz, Pz, iCP1/2, cCP1/2) for P3b and frontal channels (Fz, F1/2i, F1/2c) for Novelty P3. For the omission components, we selected temporo-parietal channels (cP3/4, cP5/6, iTP7/8, iCP5/6) for oN1, fronto-central channels for oN2 (Cz, Fz, iFC1/2, cFC1/2) and for oP3-1 (Cz, iC1/2, iFC1/2), and centro-parietal channels (Cz, CPz, iC1/2, iCP1/2, cCP1/2) for oP3-2. From here on, the selected groups of channels will be referred to as regions of interest (ROI).

### Statistical analysis

Statistics combined Bayesian and frequentist tests of behavioural data and the PCA-derived component scores. JASP was used for all statistical analyses (JASP Team, 2024). Frequentist and Bayesian repeated measures ANOVAs compared the ratio of left to right button presses and the median IPI between block types (*Predictable Sound Left*, *Predictable Sound Right*, *Unpredictable Sound*). In cases where Mauchly’s test indicated that the data violated the assumption of sphericity, degrees of freedom were adjusted using the Huynh-Feldt estimates of sphericity.

Mean component scores were computed within channels of interest for each sound and sound omission component. Bayesian and frequentist two-sided paired-samples *t*-tests then compared mean component scores between the *predicted* (80%), *mispredicted* (20%) and *unpredictable* (50%) *sound* trials (80% vs. 20%, 80% vs. 50%, 20% vs. 50%) for the sound PCA components. Similarly, Bayesian and frequentist two-sided paired-samples *t*-tests compared between the *predicted* (80%), *mispredicted* (20%) and *unpredictable* (50%) *sound omission* trials for the omission PCA component scores. To control the family-wise error rate in the multiple comparisons in the sound and sound omission PCA, all frequentist *t*-test underwent Bonferroni-Holm correction for a family of three (Holm, 1979). In Bayesian statistics, the null hypothesis assumed a standardized effect size of δ = 0, while the alternative hypothesis employed a Cauchy prior distribution cantered around 0. The scaling factor used was r = 0.707, reflecting the default setting for a “medium” effect size prior scaling in JASP. The standard priors for fixed effects (condition) and random effects (participant variability) were set to r = 0.5 and r = 1 for the Bayesian repeated measures ANOVA (Rouder et al., 2017). Bayes Factors (BF_10_) were interpreted according to the criteria of M. D. Lee & Wagenmakers, 2014.

## 3. Results

### Behavioural results

Participants successfully pressed the left and right buttons with similar frequency and pace across all block types (**Table 1**). The ratio of left to right button presses did not differ significantly between block types (*BF_10_* = 0.336, *F*(1.8, 72.0) = 1.518, *p* = .227), nor did the median IPIs (*BF_10_* = 0.344, *F*(1.75, 69.84) = 1.83, *p* = .172).

**Table 1.** shows the descriptive statistics for the ratios of button presses followed by a sound to all button presses for each block type. Below are the descriptive statistics for the inter-press intervals in milliseconds for each block type. BPR = Button Press Ratio; IPI = Inter-Press Interval in ms; PSL = Predictable Sound Left; PSR = Predictable Sound Right; US = Unpredictable Sound; *M* = mean; *SD* = standard deviation.

| Contrast | Condition | Median | *M* | *SD* | Range |
| --- | --- | --- | --- | --- | --- |
| BPR | PSL | 1.022 | 1.048 | 0.185 | 0.681 – 1.801 |
|  | PSR | 0.996 | 0.992 | 0.164 | 0.673 – 1.3.63 |
|  | US | 1.000 | 1.006 | 0.064 | 0.900 – 1.184 |
| IPI (ms) | PSL | 3828 | 3862 | 284 | 3422 – 4622 |
|  | PSR | 3816 | 3888 | 312 | 3430 – 4624 |
|  | US | 3848 | 3935 | 356 | 3365 – 4805 |

### Sound Responses

Grand-average ERPs and difference waves over the fronto-central ROI (**Figure 3**) revealed an early negativity of similar amplitude for both *mispredicted* and *unpredictable sounds*, followed by a later positivity that differed between these sound events. The early negativity is consistent with the mismatch negativity (MMN), while the later positivity presumably reflected a P300 complex. The subsequent sound-related PCA extracted 20 components, explaining 99.8% of the variance. Components were categorized as reflecting MMN, P3a, P3b, and Novelty P3 based on their topographies and latencies (**Figure 4**). Component scores for *mispredicted* (20%), *unpredictable* (50%) and *predicted* (80%) *sound* trials were compared to test for prediction-related differences. Statistical results confirmed equivalent MMN and P3a responses between *mispredicted* and *unpredictable sounds* and confirmed significantly more positive responses in later components (P3b, Novelty P3) for *mispredicted* compared to *unpredictable sound* events (**Table 2**).

**Figure 3.**
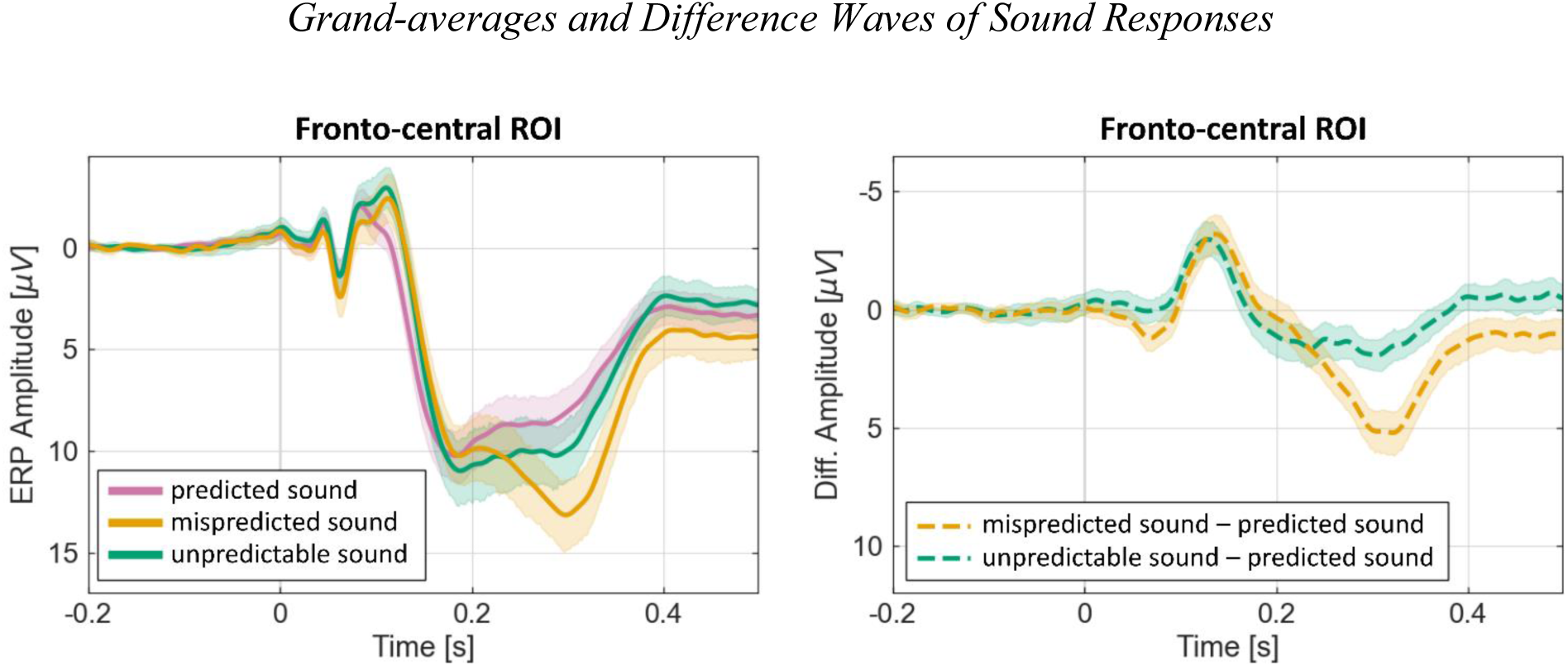
Grand-averages of predicted (80%), mispredicted (20%) and unpredictable (50%) sound responses over the fronto-central ROI (left) and difference waves over the fronto-central ROI of mispredicted minus predicted sounds (20% – 80%), and unpredictable (50% – 80%) minus predicted sounds (right). Both panels display a 95% confidence interval in transparent colours. ROI = region of interest

**Figure 4.**
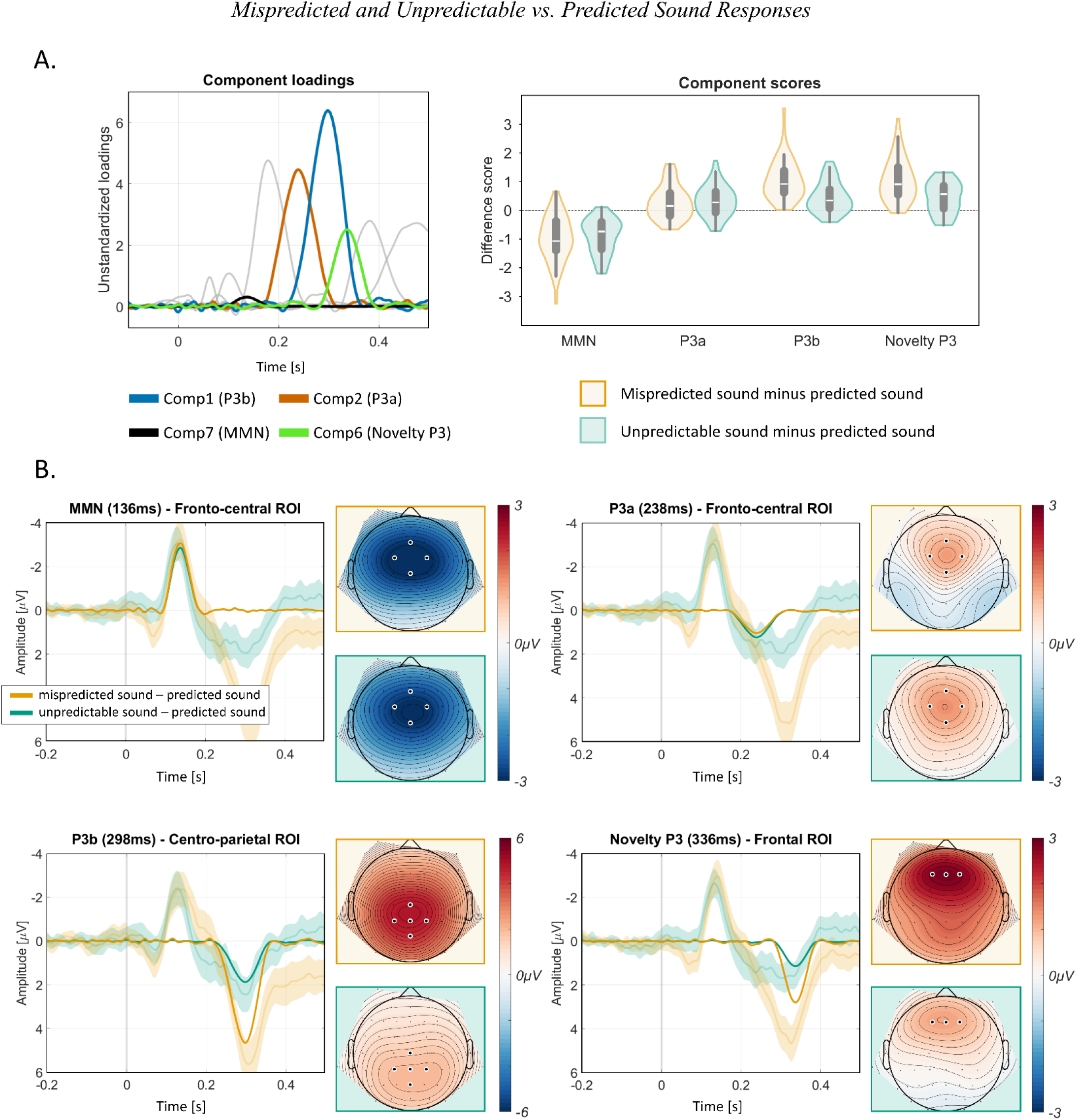
Sound PCA component loadings and difference scores for components of interest. Difference waves of sound PCA ponents (MMN, P3a, P3b, Novelty P3) with difference waves of grand average ERPs (including 95% confidence interval) ansparent colours and corresponding difference component topographies. ROI = region of interest **Panel A:** The left panel shows the sound stimulus PCA component loadings for the components of interest (MMN, P3a, P3b, elty P3). Non-relevant component loadings are displayed in grey colour. The right panel shows the difference component es for mispredicted (20%) minus predicted sound (80%) and unpredictable (50%) minus predicted (80%) sound for ponents MMN, P3a, P3b and Novelty P3 at fronto-central (MMN and P3a), centro-parietal (P3b) and frontal (Novelty P3) s. **Panel B:** Difference waves (mispredicted (20%) sound minus predicted (80%) sound, unpredictable (50%) sound minus icted (80%) sound) of sound PCA components over the regions of interest in opaque colours. Transparent colours esent grand average ERP difference waves including a 95% confidence interval. Component topographies on the right of h plot show the difference between mispredicted (20%) sound minus predicted (80%) sound conditions and between edictable (50%) minus predicted (80%) sound conditions at the latency shown.

**Table 2.** shows the results of both frequentist and Bayesian paired samples *t*-tests on the uncorrected PCA sound component scores of interest, including component 7 (MMN), component 2 (P3a), component 1 (P3b), and component 6 (Novelty P3). mispredicted = mispredicted sound trials; unpredictable = unpredictable sound trials; predicted = predicted sound trials

| Component<br>(peak latency) | Contrast | $t(40)$ | $p_{Holm}$ | Cohen's $d$ | $BF_{10}$ |
| --- | --- | --- | --- | --- | --- |
| MMN<br>(136 ms) | mispredicted vs. predicted | -7.885 | < .001 | -1.231 | 1.010*10<sup>7</sup> |
|  | unpredictable vs. predicted | -9.506 | < .001 | -1.485 | 1.111*10<sup>9</sup> |
|  | mispredicted vs. unpredictable | -0.707 | .484 | 0.110 | 0.213 |
| P3a<br>(238 ms) | mispredicted vs. predicted | 2.978 | .010 | 0.465 | 7.454 |
|  | unpredictable vs. predicted | 3.756 | < .001 | 0.587 | 51.652 |
|  | mispredicted vs. unpredictable | -0.639 | .526 | -0.100 | 0.204 |
| P3b<br>(298 ms) | mispredicted vs. predicted | 10.024 | < .001 | 1.565 | 4.708*10<sup>9</sup> |
|  | unpredictable vs. predicted | 5.297 | < .001 | 0.827 | 4121.168 |
|  | mispredicted vs. unpredictable | 6.070 | < .001 | 0.948 | 42268.951 |
| Novelty P3<br>(336 ms) | mispredicted vs. predicted | 9.126 | < .001 | 1.425 | 3.776*10<sup>8</sup> |
|  | unpredictable vs. predicted | 5.027 | < .001 | 0.785 | 1853.471 |
|  | mispredicted vs. unpredictable | 5.548 | < .001 | 0.866 | 8729.815 |

### MMN

The MMN is commonly derived from the difference between deviant and standard stimulus responses. Here, standard trials correspond to the *predicted* (80%) *sound* trials, and *mispredicted* (20%) *sound* trials correspond to deviants. In addition, we tested for the presence of an MMN component in *unpredictable* (50%) *sound* trials, relative to *predicted* (80%) *sound* trials. As shown in Figure 5, component 7 had a peak latency of 136 ms and a fronto-central scalp distribution with a negative polarity. The component explained 5.7% of variance. Its topography, polarity, and latency were compatible with an MMN. Compared to the *predicted sound* trials, both *mispredicted sound* (*BF_10_* = 1.010*10^7^) and *unpredictable sound* (*BF_10_* = 1.111*10^9^) resulted in larger amplitudes of the MMN component (stronger negativity), while *mispredicted sound* trials did not differ from *unpredictable sound* trials (*BF_10_* = 0.213).

**Figure 5.**
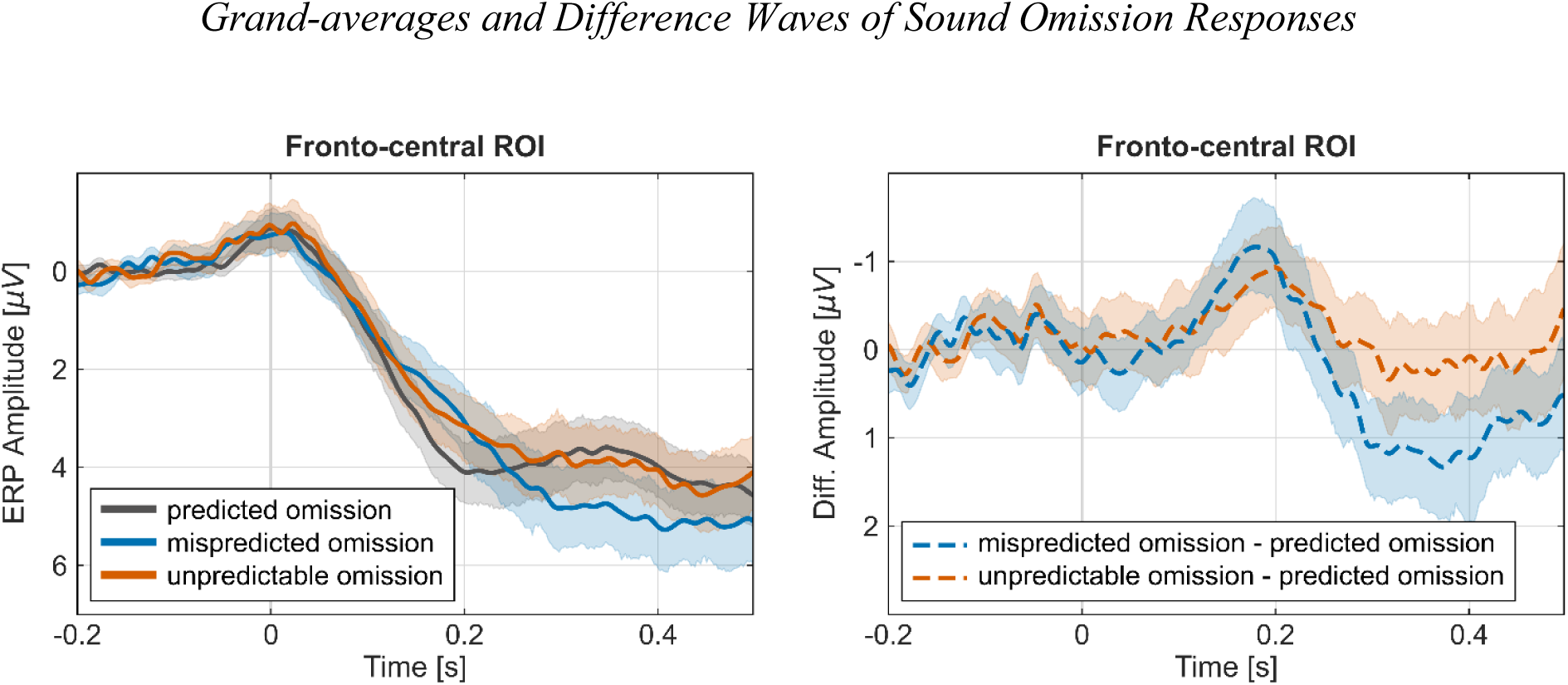
Grand-averages of predicted (80%), mispredicted (20%) and unpredictable (50%) sound omission responses over the fronto-central ROI (left) and difference waves over the fronto-central ROI of mispredicted minus predicted omissions (20% – 80%), and unpredictable (50% – 80%) minus predicted omissions (right). Both panels display a 95% confidence interval in transparent colours; ROI = region of interest

### P3a

Like the MMN, the P3a component is typically derived from the difference between deviant and standard stimulus responses. We therefore followed a similar approach as for the MMN, and compared *predicted* (80%; i.e., the standard) *sound* trials to *mispredicted* (20%) and *unpredictable* (50%) *sound* trials. As shown in Figure 5, component 2 had a peak latency of 238 ms and a fronto-central scalp distribution with a positive polarity, resembling typical features of a P3a. The component explained 17.4% of variance. The data provides moderate to strong evidence for more negative amplitudes elicited by both the *mispredicted* (*BF_10_* = 7.454) and the *unpredictable sound* trials (*BF_10_* = 51.652) in comparison to the *predicted sound* trials. Furthermore, the analysis reveals moderate evidence for the null model, i.e., no difference between *mispredicted sound* trials and *unpredictable sound* trials (*BF_10_* = 0.204)

### P3b

Component 1 captured a component with more positive amplitudes in response to *mispredicted* (20%) and *unpredictable* (50%) compared to *predicted* (80%) *sounds*, presumably reflecting P3b with a centro-parietal distribution and a peak latency of 298 ms. The component explained 22.2% of variance. The data provides extreme evidence for the alternative model, indicating a more positive amplitude for *mispredicted sound* trials vs. *predicted sound* trials (*BF_10_* = 4.708*10^9^), *unpredictable sound* trials vs. *predicted sound* trials (*BF_10_* = 4121.168) and *mispredicted sound* trials vs. *unpredictable sound* trials (*BF_10_* = 42268.951).

### Novelty P3

Component 6 captured a component with more positive amplitudes in response to *mispredicted* (20%) and *unpredictable* (50%) compared to *predicted* (80%) *sounds*, presumably reflecting Novelty P3 with a frontal distribution and had a peak latency of 336 ms. The component explained 7.1% of variance. The data provides extreme evidence for the alternative model, indicating a more positive amplitude for *mispredicted sound* trials vs. *predicted sound* trials (*BF_10_* = 3.776*10^8^) *unpredictable sound* trials vs. *predicted sound* trials (*BF_10_*= 1853.471) and *mispredicted sound* trials vs. *unpredictable sound* trial (*BF_10_* = 8729.815).

In sum, *mispredicted* (20%) and *unpredictable* (50%) *sounds* evoked increased amplitudes compared to *predicted* (80%) *sounds* reflected by the ERP-components MMN, P3a, P3b, and Novelty P3. While there was evidence for similar amplitudes of MMN and P3a in response to *mispredicted* and *unpredictable sounds*, amplitudes of P3b and Novelty P3 were increased in response to *mispredicted* compared to *unpredictable sounds*.

### Sound Omission Responses

Similar to the sound responses, grand-mean ERPs and difference waves at fronto-central channels of interest show an early negativity after sound omission presentation followed by a positivity (**Figure 5**). The omission PCA extracted a total of 18 components (as determined by EKC), explaining a total of 97.6% of variance. Component loadings and difference scores for the components capturing oN1, oN2, oP3-1, and oP3-2, in *mispredicted* (20% omission probability) *omission trials* and *unpredictable* (50%) *omission trials*, each relative to *predicted* (80%) *omission trials* were categorised based on their topographies and latencies (**Figure 6**).

**Figure 6.**
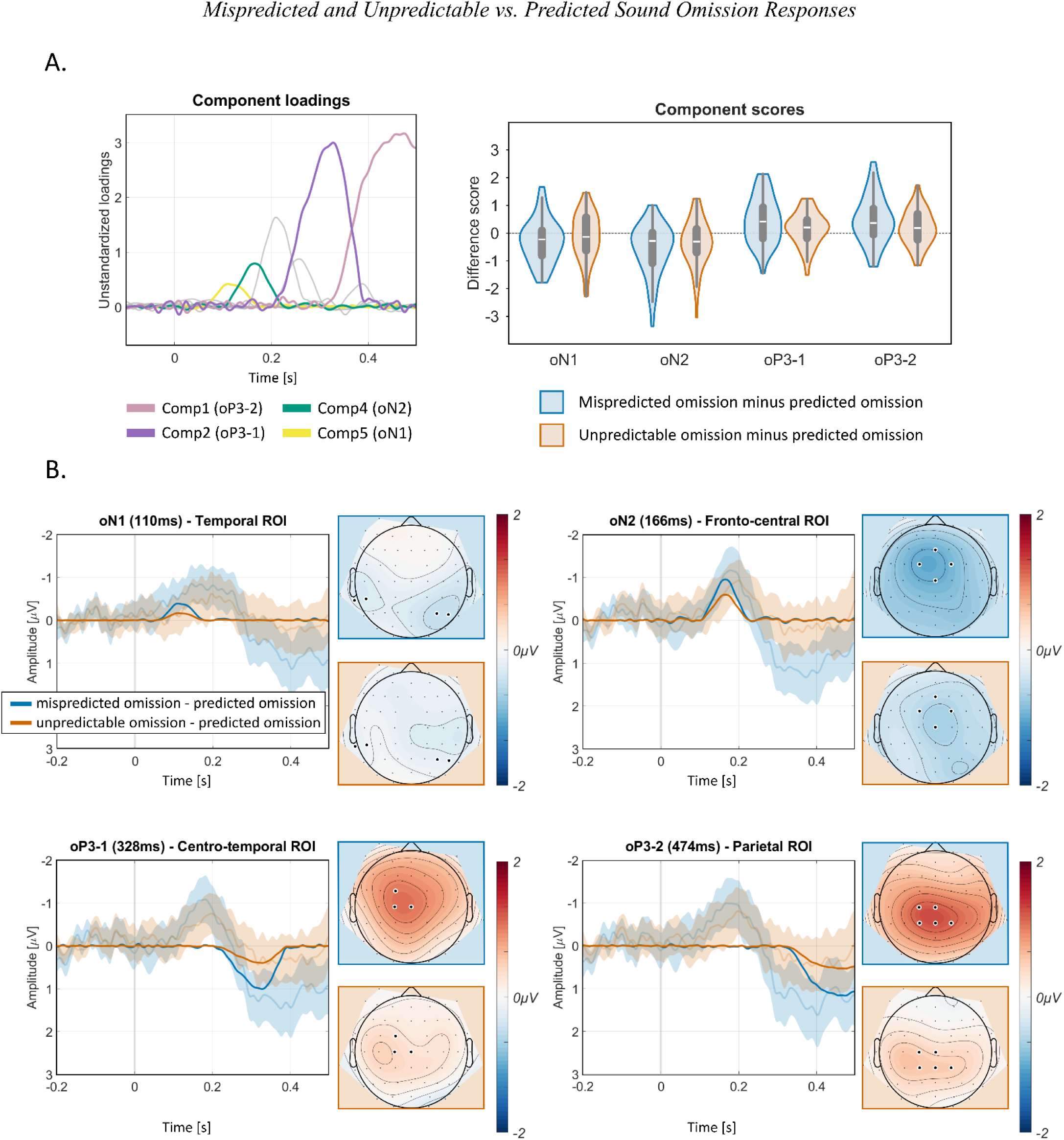
Sound omission PCA component loadings and difference scores for components of interest. Difference waves of nd omission PCA components (oN1, oN2, oP3-1, oP3-2) with difference waves of grand average ERPs (including 95% confidence interval) in transparent colours and corresponding difference component topographies. ROI = region of interest **Panel A:** The left panel shows the omission PCA component loadings for the components of interest oN1, oN2, oP3-1, oP3-2. Non-relevant component loadings are displayed in grey colour. The right panel shows the component scores for mispredicted %) minus predicted omission (80%) and unpredictable (50%) minus predicted (80%) omission for components oN1, oN2, op3-1 and oP3-2 at temporal (oN1), centro-frontal (oN2), centro-temporal (oP3-1) and parietal (oP3-2) ROIs. **Panel B:** Difference waves (mispredicted (20%) sound omission minus predicted (80%) sound omission, unpredictable (50%) nd omission minus predicted (80%) sound omission) of sound omission PCA components over the regions of interest in que colours. Transparent colours represent grand average ERP difference waves including a 95% confidence interval. ponent topographies on the right of each plot show the difference between mispredicted (20%) minus predicted (80%) nd omission trials, and between unpredictable (50%) minus predicted (80%) sound omission trials, at the latency shown.

Component scores for *mispredicted* (20%), *unpredictable* (50%) and *predicted* (80%) *omission* trials were compared to test for prediction-related differences (see **Table 3** for statistical results). While we found convincing evidence that some of the omission responses differed between *mispredicted* (20%) *omissions* and *predicted* (80%) *omissions* (**Table 3**), evidence for differences between *unpredictable* (50%) and *predicted* (80%) *omissions* and between *mispredicted* (20%) and *unpredictable* (50%) *omissions* was anecdotal or moderate at most and difficult to interpret for all omission responses. In the following paragraphs, we therefore focus on differences between *mispredicted* and *predicted omissions*.

**Table 3.** shows the results of both frequentist and Bayesian paired samples *t*-tests on the PCA omission component scores of interest, including component 5 (oN1), component 4 (oN2), component 2 (oP3-1), and component 1 (oP3-2). Mispredicted = mispredicted omission trials, predicted = predicted omission trials, unpredictable = unpredictable omission trials

| Component<br>(peak latency) | Contrast | $t(40)$ | $p_{Holm}$ | Cohen's $d$ | $BF_{10}$ |
| --- | --- | --- | --- | --- | --- |
| oN1<br>(110 ms) | mispredicted vs. predicted | -2.335 | .075 | -0.365 | 1.888 |
|  | unpredictable vs. predicted | -0.921 | .518 | -0.144 | 0.251 |
|  | mispredicted vs. unpredictable | -1.144 | .518 | -0.179 | 0.310 |
| oN2<br>(166 ms) | mispredicted vs. predicted | -4.134 | < .001 | -0.646 | 143.624 |
|  | unpredictable vs. predicted | -2.765 | .009 | -0.432 | 4.610 |
|  | mispredicted vs. unpredictable | -1.254 | .217 | -0.196 | 0.349 |
| oP3-1<br>(328 ms) | mispredicted vs. predicted | 3.475 | .003 | 0.543 | 24.985 |
|  | unpredictable vs. predicted | 1.631 | .111 | 0.255 | 0.569 |
|  | mispredicted vs. unpredictable | 2.276 | .056 | 0.355 | 1.684 |
| oP3-2<br>(474 ms) | mispredicted vs. predicted | 3.484 | .003 | 0.544 | 25.515 |
|  | unpredictable vs. predicted | 1.940 | .106 | 0.303 | 0.922 |
|  | mispredicted vs. unpredictable | 1.990 | .106 | 0.311 | 1.003 |

### oN1

Component 5, explaining 6.1% of variance, was identified as a candidate of the omission related N1 response with a mainly temporo-occipital scalp distribution and a peak latency of 110 ms. The data provides only anecdotal evidence for the alternative model, i.e., the *mispredicted omission* trials elicited a more negative amplitude than the *predicted omission* trials (*BF_10_*= 1.888).

### oN2

Component 4, explaining 7.9% of variance, has a mainly frontocentral scalp distribution and a peak latency of 166 ms. The data provides extreme evidence for the alternative model, i.e., the *mispredicted omission* trials elicited a more negative amplitude than the *predicted omission* trials (*BF_10_* = 143.624).

### oP3-1

PCA extracted two oP3 components which we labelled oP3-1 and oP3-2. Component 2 (oP3-1), explaining 22.5% of variance, has a frontocentral scalp distribution and a peak latency of 328 ms. The data provides strong evidence for the alternative model, i.e., the *mispredicted omission* trials elicited a more negative amplitude than the *predicted omission* trials (*BF_10_* = 24.985).

### oP3-2

Component 1, explaining 30.9% of variance, has a centroparietal scalp distribution and a peak latency of 474 ms. The data provides strong evidence for the alternative model, i.e., the *mispredicted omission* trials elicited a more positive amplitude than the *predicted omission* trials (*BF_10_* = 25.515).

In sum, most omission responses (oN2, oP3-1, oP3-2) showed increased amplitudes in response to *mispredicted* (20%) *omissions* compared to *predicted* (80%) *omissions*, except the oN1 component, for which we found only weak evidence for this effect.

## 4. Discussion

Predictions about the auditory effects of our actions help us determine whether our actions fulfil our intentions. However, it remains unclear whether these predictions can flexibly adapt when action-effect contingencies change frequently, as they do in real-world scenarios. We employed a dynamic button-press paradigm that required participants to simultaneously manage multiple action-sound contingencies. To clearly disentangle bottom-up and top-down effects on brain prediction error signals, we violated action-based predictions by the presentation of unexpected sounds and unexpected omission of sounds. Both evoked a series of typical error-related brain responses in the event-related potential at different levels of auditory processing: MMN and P3 complex (sounds) and oN1, oN2, oP3 complex (omissions). This demonstrates the brain’s ability to adjust action-based predictions dynamically across the hierarchical structures of the auditory processing system. Moreover, we observed similar sound-related MMN in the conditions predicting silence and the unpredictable condition, while P3 components differed. This unexpected finding will be discussed as a possible inability of the brain to reliably predict silence at the early sensory level.

### Flexible Top-down Control of Action-based Predictions of Sounds and Omissions

Self-initiated actions that violated top-down predictions evoked distinct prediction error-related components. For *mispredicted sounds* (where silence was expected but a sound presented), these included MMN, P3a, P3b, and Novelty P3. For *mispredicted omissions* (where a sound was expected but omitted), components included oN1 (with weak evidence), oN2, oP3-1, and oP3-2. These findings add further evidence that self-initiated actions cause the generation of predictions on sensory action effects (Darriba et al., 2021; Korka et al., 2019; Timm et al., 2014). Importantly, our results show that the generation of top-down predictions through actions is highly flexible and can be observed even when participants frequently switch between actions. This flexibility demonstrates that prediction generation mechanisms can dynamically switch between different effector-specific (i.e. hand specific) action-effect expectations based on the agent’s current goals. Moreover, our findings align with the eAERS (Korka et al., 2022), supporting the idea that action-effect expectations from the motor system are selectively retrieved and transmitted to the sensory predictive model in auditory processing (Hommel, 2019; Hommel et al., 2001). On the sensory level, both sound and omission error responses (MMN, oN1 and oN2) to mispredicted events are evoked in the present study. This suggests that the predictive model dynamically adapts comparative processes between sensory representations and action-effect expectations to the current task. Thus, we conclude that the retrieval and transmission of top-down information from motor systems to sensory predictive models occur flexibly. These findings corroborate the close and dynamic interplay between the motor system and auditory perception processing. Later components, such as P3b and Novelty P3 for *mispredicted sounds*, and oP3-1 and oP3-2 for *mispredicted omissions*, are suggested to reflect the evaluation of discrepancies and integration of contextual information (Polich, 2007) to form a comprehensive event representation (Korka et al., 2022). It is assumed that this representation is then used by higher-level processes to adapt attention or motor behaviour toward unexpected events (Korka et al., 2022; Ullsperger et al., 2014). As with early error responses, we found that later P3 components are evoked for *mispredicted sound* and *omission* events. This indicates a similarly high flexibility with regard to the current agent’s task (Mars et al., 2008). These results highlight the flexibility of higher-level cognitive systems (e.g., attention allocation) in response to violation of predictions.

In sum, our results provide strong evidence supporting the hypothesis that top-down predictions can be flexibly generated on a trial-by-trial basis. A violation of these predictions evoked error-signals in response to unexpected sounds and unexpected omissions on several levels of auditory processing. We suggest that auditory prediction is a highly flexible process within the hierarchically structured eAERS model, involving a close and dynamic interplay between high-level motor systems and auditory perception.

### Predictions of Silence in the Auditory Processing Hierarchy

We found that the MMN to *mispredicted sounds* (prediction of silence) was equivalent to the MMN to *unpredictable sounds* (50% *Unpredictable Sound* condition). This finding is surprising because we expected that the *mispredicted sounds* violated a prediction of silence (80% *predicted omission* trials), while no reliable prediction could be formed in the *Unpredictable Sound* condition. According to the eAERS (Korka et al., 2022), the MMN responses are related to the early low-level comparison of sensory input to top-down predictions. This comparison appears straightforward for actual auditory stimuli, such as sounds following a button press. In this case, the sensory input has basic acoustic features (e.g., intensity, frequency) which, together with its acoustic context (e.g., such as sound patterns), can be the content of top-down predictions. Resulting prediction errors and subsequent updating of the predictive model cause the observed MMN effects (István Winkler, 2007). However, the representation of silence as an anticipated event is less straightforward. There are no actual features of silence a top-down auditory prediction could anticipate, and it is therefore unclear how the prediction of silence—the absence of features—would be represented neurally (in a network dedicated to the processing of sound features). Furthermore, unlike silence, sound in our study corresponded to a change in sensory input at a discrete point in time. Thus, the comparative processes that involve top-down predictions of expected silence, in particular as a continuous event without change in sensory input, pose a challenge in predictive coding.

On this notion, the present results reveal evidence for an equally strong activation of this early error detection system to different kinds of sound presentations: in a predictable condition when silence followed a button press in the majority of times (expected silence) and in an unpredictable condition when it was equally likely that a sound or an omission followed a button press (no clear expectation). **Figure 7** provides a visualization of how prediction errors manifest under these conditions. In the predictable condition, predictions for sounds or omissions are formed based on the most probable motor action effects, and prediction errors arise when sensory input mismatches these predictions. In contrast, in the unpredictable condition, no reliable sensory-level predictions can be formed due to the 50% probability of sound or omission. Here, the equivalent MMN responses we observed for *mispredicted sounds* and *unpredictable sounds* demonstrate, that despite the clear predictability of silence, an equally strong error response is triggered as in the unpredictable condition. Thus, if the prediction error for *mispredicted sounds* (silence predicted) and *unpredictable sounds* (i.e., *no* prediction possible) is equivalent, we can conclude that also the predictions are equivalent. That is, apparently encoding silence as a reliable prediction at sensory levels is not possible. Therefore, the MMN observed in this study in response to *mispredicted* and *unpredictable sounds* can be interpreted as a prediction error at the sensory level inherent to the exogenous sound response in the absence of a prediction (Schröger et al., 2015). This MMN-related prediction error is revealed by subtracting the ERP of a *predicted sound* that does not elicit a prediction error response. In contrast, later P3 components (P3b, Novelty P3) reflecting conceptual processing reveal significant differences between the violation of expected silence (80% *predicted omission*) by sounds and the unpredictable condition (50%). According to the eAERS, responses at this stage represent the evaluation of resulting discrepancies and the integration of additional contextual information outside the auditory environment (i.e., motor action) with the sensory representation to form a comprehensive auditory event representation (Polich, 2007). This representation is further fed into a higher-level event predictive model which contains an abstract representation of the expected auditory event including its non-auditory context for further comparative processes (Korka et al., 2022). The significantly increased amplitudes of P3b and Novelty P3 components to unexpected sounds in the 80% predicted omission condition compared to the unpredictable condition indicate, that expected silence is only adequately represented when additional contextual information (i.e., prior motor action or task requirements) has been integrated. This integration at later response stages underscores the hierarchical nature of auditory processing, where silence is represented conceptually only at higher cognitive levels within the eAERS framework.

**Figure 7.**
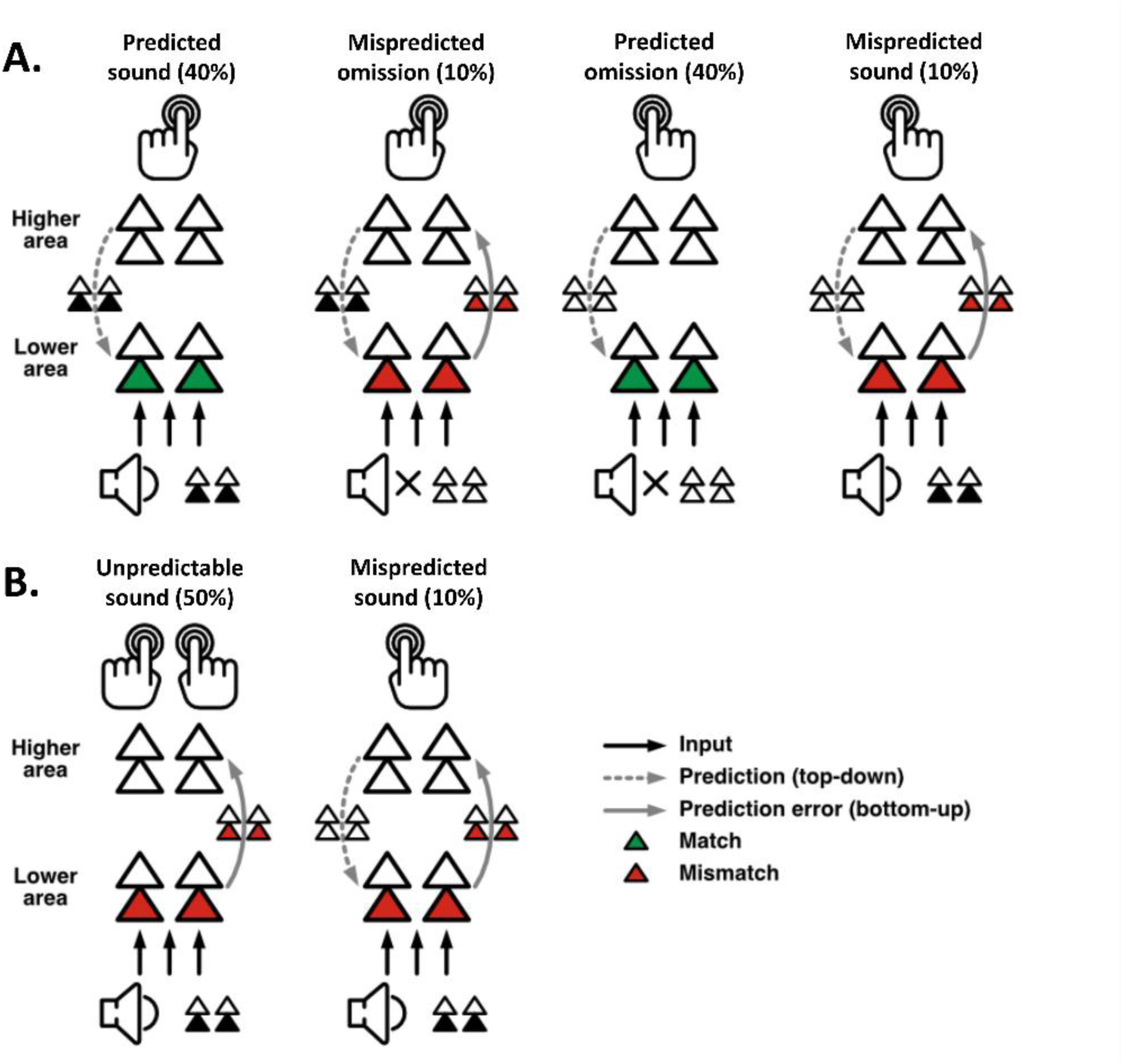
Schematic description of action-sound predictions. **Panel A:** Prediction error in the predictable condition (based on Schröger et al., 2015, Fig. 2, adapted): Depending on the button press a sound (left hand) or no sound (right hand) is expected. The expectation is transferred top-down as prediction to sensory areas. In case the sensory input mismatches the prediction, the prediction error is transferred bottom-up to higher areas to update the generative model. The observed prediction error responses (oN1 and oN2 for mispredicted omissions and MMN for mispredicted sounds) demonstrate that predictions can be established at sensory levels on a trial-by-trial basis. **Panel B:** Prediction error for unpredictable vs. mispredicted sounds: In the unpredictable condition, no reliable top-down prediction can be established at sensory levels. We observed equivalent prediction error responses for unpredictable sounds and mispredicted sounds (in a strict sense, there can be no prediction error without prediction, which is why Schröger et al., 2015 also refer to it as “exogenous response” in the former case). We did not see an additional ERP signature of expectation violation for predicted silence by sounds compared to no prediction by sounds. Our interpretation of this surprising finding is that the representation of the prediction of an expected omission (i.e. silence) at sensory levels is equivalent to the absence of a prediction. We consider the possible alternative interpretation that omissions (silence) are consistently predicted in the unpredictable condition as implausible because sounds were frequently encountered.

On a broader level of implication, auditory information, omnipresent and complex, constitutes a fundamental aspect of our environment. The brain’s proficiency in processing a diverse array of auditory cues at pre-attentive stages, reflected by mechanisms such as the MMN (see Fitzgerald & Todd, 2020; Kraus & Chandrasekaran, 2010; Näätänen et al., 2007 for a comprehensive overview) underlines its fine-tuned sensitivity to sound perception. Given that the brain is inherently designed to react to stimuli, the absence of stimuli (silence) may therefore not be encoded in the same way as sound. Traditionally, silence as an auditory event is considered to be derived from the absence of sound (Bregman & McAdams, 1994; Kubovy & van Valkenburg, 2001; István Winkler et al., 2009). However, in a recent study conducted by Goh et al. (2023), silence is described as not just the absence of sound but as a sound-like entity, understanding silence to be an active element in the auditory environment. In a series of experiments, they showed that silences can substitute for sounds in event-based auditory illusions. Their findings show that auditory perception handles moments of silence similarly to sounds at a behavioural level. One explanation by the authors, and in line with TEC (Hommel, 2019), is that similar to perceived sound, perceived silence can be understood as an empty event file (Hommel, 2004; Hommel et al., 2001). This means that, despite the lack of explicit auditory cues, silence carries non-acoustic sensory information, such as temporal and contextual properties. According to the differences we found in later P3 components, the present results align well with this interpretation of silence as an “empty” auditory event file and the eAERS. It appears that an explicit representation can only be achieved when relevant non-auditory information has been integrated. This leads to the conclusion that a representation of silence is not solely derived from the absence of sound. However, due to the lack of actual auditory information at lower sensory levels, silence can only be perceived as an auditory event when relevant non-auditory information has been integrated at higher auditory processing levels. Together, these findings show that silence, much like sound, plays an active role in our sensory environment and, based on our findings, predictions of silence can be represented at conceptual but not necessarily at sensory levels. In other words, presumably silence can be represented as a concept depending on contextual information but not as explicit sensation in a strict sense.

This interpretation challenges traditional views of silence as merely the absence of sound, supporting its conceptualization as an “empty” event file within the TEC framework. This assumption accounts for the equal error response to *mispredicted sounds* when silence is expected and *unpredictable sounds* at early response stages on the one hand, as depicted in **Figure 4**. On the other hand, the significantly different magnitudes at later response stages demonstrate that silence can be identified as a discrete auditory event in higher cognitive stages by the integration of additional contextual information.

## 5. Limitations

The ERPs in response to sound omissions presented a unique challenge due to the nature of silence as an auditory event. Unlike sound, silence has no actual auditory features and therefore no explicit onset or offset cue. To address this, silence and sound events were coupled to a button press. These button presses acted as onset cues for the sound and the sound omission. However, the IPI of three seconds was significantly longer than typically used in other omission studies (Dercksen et al., 2020; SanMiguel, Widmann, et al., 2013; Widmann & Schröger, 2022). Moreover, the IPI consisted of silence only. In our study, this extended silent gap is combined with the inherent limitations of human temporal perception, which is context dependent (Ross & Balasubramaniam, 2022) and thus possibly not accurate to the millisecond. Together, this likely contributed to a variability in the timing of when exactly participants perceived the (predicted) silence as an action effect within the stream of silence between button presses. This variability in perception may lead to a temporal distribution of oN1 responses. As event-related potentials (ERPs) are averaged over multiple trials and participants, this distribution likely results in an attenuation or flattening of the oN1 signal, making it less pronounced compared to findings in studies with shorter IPIs.

## 6. Conclusion and Outlook

Our findings demonstrate the brain’s ability to flexibly adapt action-based auditory predictions in dynamic environments, demonstrated by prediction error signals in sound and omission ERPs. Notably, the equivalent MMN responses to mispredicted and unpredictable sounds suggest that silence may not be explicitly represented at early sensory levels, requiring higher-level contextual integration for processing. These results motivate future studies to explore the role of silence in auditory prediction and its integration into the eAERS model. For instance, how is silence as action-based prediction processed in complex auditory environments like speech or music? Such investigations could extend the understanding of sensory-motor integration and refine predictive coding frameworks. Beyond basic research, our findings have broader implications for real-world applications. The ability to dynamically adjust predictions is essential for navigating complex auditory environments, such as social interactions or noisy settings. Additionally, these insights may inform clinical research, particularly in disorders like schizophrenia or autism, where atypical responses to auditory prediction errors are common (Haigh et al., 2023; Kaser et al., 2013; M. Lee et al., 2017). Understanding how these populations process silence and unexpected sounds could lead to novel diagnostic or therapeutic approaches. In summary, this study highlights the flexibility of action-based auditory predictions and raises important questions about the neural representation of silence, paving the way for future research and practical applications.

## Author Contributions

**Fabian Aurich:** Conceptualization, Data curation, Validation, Formal analysis, Investigation, Writing – original draft, Writing – review and editing, Visualization. **Andreas Widmann:** Conceptualization, Methodology, Software, Writing – original draft, Writing – review and editing, Visualization. **Tjerk T. Dercksen:** Software, Validation, Formal analysis, Data curation, Writing – review and editing. **Betina Korka:** Conceptualization, Methodology, Writing – review and editing. **Anni Richter:** Conceptualization, Writing – review and editing. **Max-Philipp Stenner:** Conceptualization, Methodology, Validation, Resources, Writing – review and editing, Supervision, Project administration, Funding acquisition. **Nicole Wetzel:** Conceptualization, Methodology, Validation, Resources, Writing – review and editing, Supervision, Project administration, Funding acquisition.

## Data Availability Statement

The data supporting the results of this study are available from the corresponding author upon request.

## Acknowledgements

The authors thank all participants that took part in this study. The authors would also like to thank Dunja Kunke, Gabriele Schöps and Marit Giechau for support in data acquisition.

M.-P.S. was supported by a VolkswagenStiftung Freigeist Fellowship, project-ID 92977, and received funding from a Deutsche Forschungsgemeinschaft Sonderforschungsbereich, SFB-1436, TPC03, project-ID 425899996.

N.W. was supported by the Center for Behavioral Brain Sciences Magdeburg financed by the European Regional Development Fund (ZS/2016/04/78120) and Leibniz Association (P58/2017) and received funding from a DFG (WE5026/4-1).

## Supplementary

**Table A1:**
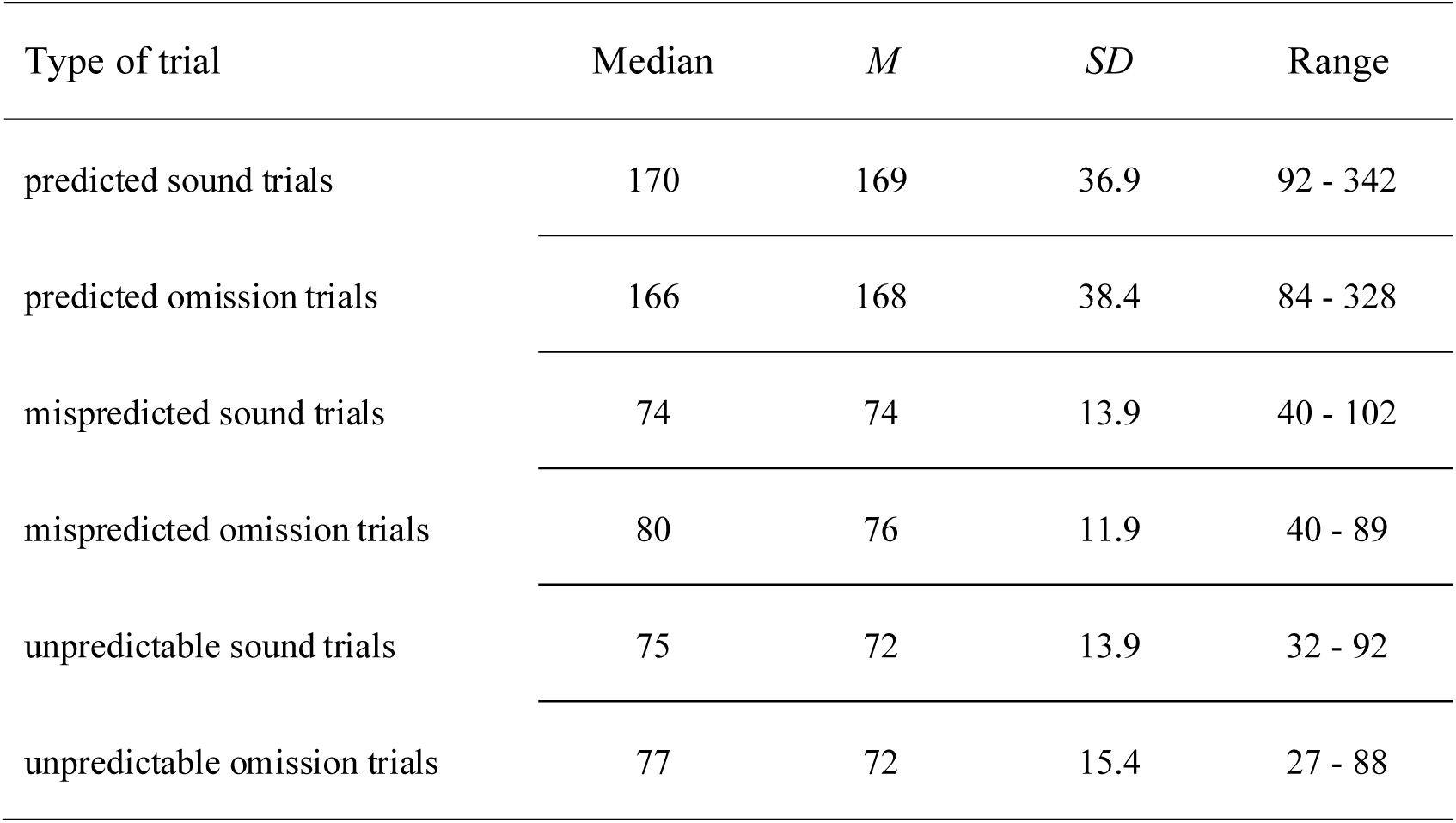
Descriptive statistics for included trials per trial type. *M = mean, SD = standard deviation*

## References

1. Arnal, L. H., & Giraud, A.-L. (2012). Cortical oscillations and sensory predictions. Trends in Cognitive Sciences, 16(7), 390–398. 10.1016/j.tics.2012.05.003

2. Baragona, V., Schröger, E., & Widmann, A. (2025). Salient, Unexpected Omissions of Sounds Can Involuntarily Distract Attention. Journal of Cognitive Neuroscience, 1–16. 10.1162/jocn_a_02307

3. Barry, R. J., & Rushby, J. A. (2006). An orienting reflex perspective on anteriorisation of the P3 of the event-related potential. Experimental Brain Research, 173(3), 539–545. 10.1007/s00221-006-0590-8

4. Barry, R. J., Steiner, G. Z., Blasio, F. M. de, Fogarty, J. S., Karamacoska, D., & MacDonald, B. (2020). Components in the P300: Don’t forget the Novelty P3! Psychophysiology, 57(7), e13371. 10.1111/psyp.13371

5. Bigdely-Shamlo, N., Mullen, T., Kothe, C., Su, K.-M., & Robbins, K. A. (2015). The PREP pipeline: Standardized preprocessing for large-scale EEG analysis. Frontiers in Neuroinformatics, 9, 16. 10.3389/fninf.2015.00016

6. Braeken, J., & van Assen, M. A. L. M. (2017). An empirical Kaiser criterion. Psychological Methods, 22(3), 450–466. 10.1037/met0000074

7. Brainard, D. H. (1997). The Psychophysics Toolbox. Spatial Vision, 10(4), 433–436.

8. Bregman, A. S., & McAdams, S. (1994). Auditory Scene Analysis: The Perceptual Organization of Sound. The Journal of the Acoustical Society of America, 95(2), 1177–1178. 10.1121/1.408434

9. Clark, A. (2013). Whatever next? Predictive brains, situated agents, and the future of cognitive science. The Behavioral and Brain Sciences, 36(3), 181–204. 10.1017/S0140525X12000477

10. Darriba, Á., Hsu, Y.-F., van Ommen, S., & Waszak, F. (2021). Intention-based and sensory-based predictions. Scientific Reports, 11(1), 19899. 10.1038/s41598-021-99445-z

11. Delorme, A., & Makeig, S. (2004). Eeglab: An open source toolbox for analysis of single-trial EEG dynamics including independent component analysis. Journal of Neuroscience Methods, 134(1), 9–21. 10.1016/j.jneumeth.2003.10.009

12. Dercksen, T. T., Widmann, A., Noesselt, T., & Wetzel, N. (2024). Somatosensory omissions reveal action-related predictive processing. Human Brain Mapping, 45(4), e26550. 10.1002/hbm.26550

13. Dercksen, T. T., Widmann, A., Schröger, E., & Wetzel, N. (2020). Omission related brain responses reflect specific and unspecific action-effect couplings. NeuroImage, 215, 116840. 10.1016/j.neuroimage.2020.116840

14. Elsner, B., & Hommel, B. (2001). Effect anticipation and action control. Journal of Experimental Psychology. Human Perception and Performance, 27(1), 229–240. 10.1037//0096-1523.27.1.229

15. Escera, C., Alho, K., Winkler, I., & Näätänen, R. (1998). Neural mechanisms of involuntary attention to acoustic novelty and change. Journal of Cognitive Neuroscience, 10(5), 590–604. 10.1162/089892998562997

16. Faul, F., Erdfelder, E., Buchner, A., & Lang, A.-G. (2009). Statistical power analyses using G*Power 3.1: Tests for correlation and regression analyses. Behavior Research Methods, 41(4), 1149–1160. 10.3758/BRM.41.4.1149

17. Feldman, H., & Friston, K. J. (2010). Attention, uncertainty, and free-energy. Frontiers in Human Neuroscience, 4, 215. 10.3389/fnhum.2010.00215

18. Fitzgerald, K., & Todd, J. (2020). Making Sense of Mismatch Negativity. Frontiers in Psychiatry, 11, 468. 10.3389/fpsyt.2020.00468

19. Friston, K. (2010). The free-energy principle: A unified brain theory? Nature Reviews. Neuroscience, 11(2), 127–138. 10.1038/nrn2787

20. Garrido, M. I., Kilner, J. M., Stephan, K. E., & Friston, K. J. (2009). The mismatch negativity: A review of underlying mechanisms. Clinical Neurophysiology : Official Journal of the International Federation of Clinical Neurophysiology, 120(3), 453–463. 10.1016/j.clinph.2008.11.029

21. Goh, R. Z., Phillips, I. B., & Firestone, C. (2023). The perception of silence. Proceedings of the National Academy of Sciences of the United States of America, 120(29), e2301463120. 10.1073/pnas.2301463120

22. Greenwald, A. G. (1970). Sensory feedback mechanisms in performance control: With special reference to the ideo-motor mechanism (Vol. 77). https://psycnet.apa.org/record/1970-05997-001 10.1037/h0028689

23. Haigh, S. M., van Key, L., Brosseau, P., Eack, S. M., Leitman, D. I., Salisbury, D. F., & Behrmann, M. (2023). Assessing Trial-to-Trial Variability in Auditory ERPs in Autism and Schizophrenia. Journal of Autism and Developmental Disorders, 53(12), 4856–4871. 10.1007/s10803-022-05771-0

24. Holm, S. (1979). A Simple Sequentially Rejective Multiple Test Procedure. Scandinavian Journal of Statistics, 6(2), 65–70. http://www.jstor.org/stable/4615733

25. Hommel, B. (2003). Acquisition and control of voluntary action. In Voluntary action: Brains, minds, and sociality (pp. 34–48). Oxford University Press.

26. Hommel, B. (2004). Event files: Feature binding in and across perception and action. Trends in Cognitive Sciences, 8(11), 494–500. 10.1016/j.tics.2004.08.007

27. Hommel, B.. (2013). Ideomotor Action Control: On the Perceptual Grounding of Voluntary Actions and Agents. In W. Prinz, M. Beisert, & A. Herwig (Eds.), Action science: Foundations of an emerging discipline (pp. 113–136). MIT Press. 10.7551/mitpress/9780262018555.003.0008

28. Hommel, B. (2019). Theory of Event Coding (TEC) V2.0: Representing and controlling perception and action. Attention, Perception & Psychophysics, 81(7), 2139–2154. 10.3758/s13414-019-01779-4

29. Hommel, B., Müsseler, J., Aschersleben, G., & Prinz, W. (2001). The Theory of Event Coding (TEC): A Framework for Perception and Action Planning. The Behavioral and Brain Sciences, 24(5), 849–78; discussion 878-937. 10.1017/s0140525x01000103

30. Horváth, J., Winkler, I [István], & Bendixen, A. (2008). Do N1/MMN, P3a, and RON form a strongly coupled chain reflecting the three stages of auditory distraction? Biological Psychology, 79(2), 139–147. 10.1016/j.biopsycho.2008.04.001

31. JASP Team. (2024). JASP (Version 0.19.0)[Computer software]. https://jasp-stats.org/

32. Kaser, M., Soltesz, F., Lawrence, P., Miller, S., Dodds, C., Croft, R., Dudas, R. B., Zaman, R., Fernandez-Egea, E., Müller, U., Dean, A., Bullmore, E. T., & Nathan, P. J. (2013). Oscillatory underpinnings of mismatch negativity and their relationship with cognitive function in patients with schizophrenia. PloS One, 8(12), e83255. 10.1371/journal.pone.0083255

33. Klug, M., & Gramann, K. (2021). Identifying key factors for improving ICA-based decomposition of EEG data in mobile and stationary experiments. The European Journal of Neuroscience, 54(12), 8406–8420. 10.1111/ejn.14992

34. Korka, B., Schröger, E., & Widmann, A. (2019). Action Intention-based and Stimulus Regularity-based Predictions: Same or Different? Journal of Cognitive Neuroscience, 31(12), 1917–1932. 10.1162/jocn_a_01456

35. Korka, B., Schröger, E., & Widmann, A. (2020). What exactly is missing here? The sensory processing of unpredictable omissions is modulated by the specificity of expected action-effects. The European Journal of Neuroscience, 52(12), 4667–4683. 10.1111/ejn.14899

36. Korka, B., Widmann, A., Waszak, F., Darriba, Á., & Schröger, E. (2022). The auditory brain in action: Intention determines predictive processing in the auditory system-A review of current paradigms and findings. Psychonomic Bulletin & Review, 29(2), 321–342. 10.3758/s13423-021-01992-z

37. Kraus, N., & Chandrasekaran, B. (2010). Music training for the development of auditory skills. Nature Reviews. Neuroscience, 11(8), 599–605. 10.1038/nrn2882

38. Kubovy, M., & van Valkenburg, D. (2001). Auditory and visual objects. Cognition, 80(1-2), 97–126. 10.1016/s0010-0277(00)00155-4

39. Lee, M., Sehatpour, P., Hoptman, M. J., Lakatos, P., Dias, E. C., Kantrowitz, J. T., Martinez, A. M., & Javitt, D. C. (2017). Neural mechanisms of mismatch negativity dysfunction in schizophrenia. Molecular Psychiatry, 22(11), 1585–1593. 10.1038/mp.2017.3

40. Lee, M. D., & Wagenmakers, E.-J. (2014). Bayesian cognitive modeling: A practical course. Cambridge University Press.

41. Li, X., Liang, Z., Kleiner, M., & Lu, Z.-L. (2010). Rtbox: A device for highly accurate response time measurements. Behavior Research Methods, 42(1), 212–225. 10.3758/BRM.42.1.212

42. Makeig, S., Bell, A., Jung, T.-P., & Sejnowski, T. J. (1995). Independent Component Analysis of Electroencephalographic Data. In D. Touretzky, M.C. Mozer, & M. Hasselmo (Eds.), Advances in Neural Information Processing Systems (Vol. 8). MIT Press. https://proceedings.neurips.cc/paper_files/paper/1995/file/754dda4b1ba34c6fa89716b85d68532b-Paper.pdf

43. Mars, R. B., Debener, S., Gladwin, T. E., Harrison, L. M., Haggard, P., Rothwell, J. C., & Bestmann, S. (2008). Trial-by-trial fluctuations in the event-related electroencephalogram reflect dynamic changes in the degree of surprise. The Journal of Neuroscience, 28(47), 12539–12545. 10.1523/JNEUROSCI.2925-08.2008

44. Näätänen, R. (1990). The role of attention in auditory information processing as revealed by event-related potentials and other brain measures of cognitive function. The Behavioral and Brain Sciences, 13(2), 201–233. 10.1017/S0140525X00078407

45. Näätänen, R. (2009). Somatosensory mismatch negativity: A new clinical tool for developmental neurological research? Developmental Medicine and Child Neurology, 51(12), 930–931. 10.1111/j.1469-8749.2009.03386.x

46. Näätänen, R., Paavilainen, P., Rinne, T., & Alho, K. (2007). The mismatch negativity (MMN) in basic research of central auditory processing: A review. Clinical Neurophysiology: Official Journal of the International Federation of Clinical Neurophysiology, 118(12), 2544–2590. 10.1016/j.clinph.2007.04.026

47. Nittono, H. (2006). Voluntary stimulus production enhances deviance processing in the brain. International Journal of Psychophysiology: Official Journal of the International Organization of Psychophysiology, 59(1), 15–21. 10.1016/j.ijpsycho.2005.06.008

48. Oldfield, R. C. (1971). The assessment and analysis of handedness: The Edinburgh inventory. Neuropsychologia, 9(1), 97–113. 10.1016/0028-3932(71)90067-4

49. Pazo-Alvarez, P., Cadaveira, F., & Amenedo, E. (2003). Mmn in the visual modality: A review. Biological Psychology, 63(3), 199–236. 10.1016/s0301-0511(03)00049-8

50. Pion-Tonachini, L., Kreutz-Delgado, K., & Makeig, S. (2019). Iclabel: An automated electroencephalographic independent component classifier, dataset, and website. NeuroImage, 198, 181–197. 10.1016/j.neuroimage.2019.05.026

51. Polich, J. (2007). Updating P300: An integrative theory of P3a and P3b. Clinical Neurophysiology, 118(10), 2128–2148. 10.1016/j.clinph.2007.04.019

52. Ross, J. M., & Balasubramaniam, R. (2022). Time Perception for Musical Rhythms: Sensorimotor Perspectives on Entrainment, Simulation, and Prediction. Frontiers in Integrative Neuroscience, 16, 916220. 10.3389/fnint.2022.916220

53. Rouder, J. N., Morey, R. D., Verhagen, J., Swagman, A. R., & Wagenmakers, E.-J. (2017). Bayesian analysis of factorial designs. Psychological Methods, 22(2), 304–321. 10.1037/met0000057

54. SanMiguel, I., Saupe, K., & Schröger, E. (2013). I know what is missing here: Electrophysiological prediction error signals elicited by omissions of predicted “what” but not “when”. Frontiers in Human Neuroscience, 7, 407. 10.3389/fnhum.2013.00407

55. SanMiguel, I., Widmann, A., Bendixen, A., Trujillo-Barreto, N., & Schröger, E. (2013). Hearing silences: Human auditory processing relies on preactivation of sound-specific brain activity patterns. The Journal of Neuroscience, 33(20), 8633–8639. 10.1523/JNEUROSCI.5821-12.2013

56. Scharf, F., Widmann, A., Bonmassar, C., & Wetzel, N. (2022). A tutorial on the use of temporal principal component analysis in developmental ERP research – Opportunities and challenges. Developmental Cognitive Neuroscience, 54, 101072. 10.1016/j.dcn.2022.101072

57. Schröger, E., Marzecová, A., & SanMiguel, I. (2015). Attention and prediction in human audition: a lesson from cognitive psychophysiology. European Journal of Neuroscience, 41(5), 641–664. 10.1111/ejn.12816

58. Tast, V., Schröger, E., & Widmann, A. (2024). Suppression and omission effects in auditory predictive processing—Two of the same? *European Journal of Neuroscience*, Article ejn.16393. Advance online publication. 10.1111/ejn.16393

59. Timm, J., SanMiguel, I., Keil, J., Schröger, E., & Schönwiesner, M. (2014). Motor intention determines sensory attenuation of brain responses to self-initiated sounds. Journal of Cognitive Neuroscience, 26(7), 1481–1489. 10.1162/jocn_a_00552

60. van Laarhoven, T., Stekelenburg, J. J., & Vroomen, J. (2017). Temporal and identity prediction in visual-auditory events: Electrophysiological evidence from stimulus omissions. Brain Research, 1661, 79–87. 10.1016/j.brainres.2017.02.014

61. Wacongne, C., Labyt, E., van Wassenhove, V., Bekinschtein, T., Naccache, L., & Dehaene, S. (2011). Evidence for a hierarchy of predictions and prediction errors in human cortex. Proceedings of the National Academy of Sciences of the United States of America, 108(51), 20754–20759. 10.1073/pnas.1117807108

62. Waszak, F., & Herwig, A. (2007). Effect anticipation modulates deviance processing in the brain. Brain Research, 1183, 74–82. 10.1016/j.brainres.2007.08.082

63. Widmann, A., & Schröger, E. (2022). Intention-based predictive information modulates auditory deviance processing. Frontiers in Neuroscience, 16, 995119. 10.3389/fnins.2022.995119

64. Winkler, I [Irene], Debener, S., Müller, K.-R., & Tangermann, M. (2015). On the influence of high-pass filtering on ICA-based artifact reduction in EEG-ERP. Annual International Conference of the IEEE Engineering in Medicine and Biology Society. IEEE Engineering in Medicine and Biology Society. Annual International Conference, 2015, 4101–4105. 10.1109/EMBC.2015.7319296

65. Winkler, I [István] (2007). Interpreting the Mismatch Negativity. Journal of Psychophysiology, 21(3-4), 147–163. 10.1027/0269-8803.21.34.147

66. Winkler, I [István], Denham, S. L., & Nelken, I. (2009). Modeling the auditory scene: Predictive regularity representations and perceptual objects. Trends in Cognitive Sciences, 13(12), 532–540. 10.1016/j.tics.2009.09.003

67. Winkler, I [István], & Schröger, E. (2015). Auditory perceptual objects as generative models: Setting the stage for communication by sound. Brain and Language, 148, 1–22. 10.1016/j.bandl.2015.05.003

